# Biasing RNA coarse-grained folding simulations with Small–Angle X–ray Scattering (SAXS) data

**DOI:** 10.1101/2021.03.29.437449

**Authors:** L. Mazzanti, L. Alferkh, E. Frezza, S. Pasquali

**Affiliations:** Laboratoire CiTCoM, CNRS UMR 8038, Université de Paris, 4 Avenue de l’observatoire, 75006 Paris, France

## Abstract

RNA molecules can easily adopt alternative structures in response to different environmental conditions. As a result, a molecule’s energy landscape is rough and can exhibits a multitude of deep basins. In the absence of a high-resolution structure, Small Angle X-ray Scattering data (SAXS) can narrow down the conformational space available to the molecule and be used in conjunction with physical modeling to obtain high-resolution putative structures to be further tested by experiments. Because of the low-resolution of this data, it is natural to implement the integration of SAXS data into simulations using a coarse-grained representation of the molecule, allowing for much wider searches and faster evaluation of SAXS theoretical intensity curves than with atomistic models. We present here the theoretical framework and the implementation of a simulation approach based on our coarse-grained model HiRE-RNA combined with SAXS evaluations “on-the-fly” leading the simulation toward conformations agreeing with the scattering data, starting from partially folded structures as the ones that can easily be obtained from secondary structures predictions based tools. We show on three benchmark systems how our approach can successfully achieve high-resolution structures with remarkable similarity with the native structure recovering not only the overall shape, as imposed by SAXS data, but also the details of initially missing base pairs.

## 1 Introduction

As RNA molecules are recognized more and more to play a central role in the cell’s machinery, especially at the level of gene expression and regulation [1, 2], determining experimentally their structures remains a challenge, as shown by the relatively small number of high-resolution structures available in the Nucleic acids Data Bank in comparison with available protein structures. Part of the problem lies in the fact that RNA molecules are able to adopt alternative conformations in which the molecule rearranges its secondary structure, resulting in completely different topologies in response to the environment [3, 4].

Small-Angle X-ray Scattering has become in the last decade one of the standard experiments to gather information on molecular structures at low resolution [5, 6] and it is particularly well suited to nucleic acids because, being charged molecules, they have a high contrast with the solvent, making experimental data less noisy. With the molecules in solutions, in an adjustable environment close to natural, SAXS data provide invaluable insights on the molecule’s shape and global architecture. It is sensitive to the changes induced on the molecule by the environment that can be recreated in the test tube, and it is indeed possible to follow the conformational changes of RNA molecules subject to temperature and ion concentrations variations [7, 8, 9] as well as folding, in time resolved experiments [10]. The information from SAXS data, where the complexity of an object with 3N degrees of freedom is collapsed into one single scattering curve, needs however a detailed underlying molecular model to generate structures at high-resolutions, where the details of the molecule’s behavior can be observed. This is why SAXS data has mainly been used as a filter to be applied *a posteriori* on an ensemble of candidate high-resolution structures, generated from X-ray and NMR data on sub-parts, to extract the ones that are mostly compatible with the data. This approach is based on the ability to compute analytically the scattering curve of a molecule from its atoms coordinates and to compare it with the target experimental curve [11, 12].

In the absence of high-resolution experimental structures for the molecule’s domains, computational methods can generate the ensemble of structures needed to make the selection. Since the milestones experiments in the late 90’s showing that folding of an RNA is led primarily by the formation of its secondary structure (2D) [13], computational efforts have focused on determining the 2D structure of a molecule to then generate three-dimensional (3D) structures to help fill the gap between known sequences and available high-resolution structures [14, 15]. Recently, these predictions have benefited from the integration of chemical probing data, giving more reliable secondary structures to be used to build functional 3D models [16]. Nowadays several 3D structure prediction methods exist combining physical modeling, 2D structure predictions, chemical probing data, NMR and mutations in integrative schemes [17, 18, 19, 20].

In the case of molecules for which local structures are well characterized either by experiments or by computational predictions, a filtering strategy to find structures agreeing with SAXS data can be very successful as the search in the conformational space reduces to exploring a limited number of degrees of freedom concerning the rotations of different domains or helices with respect to each other [21, 22, 23]. It is also possible to use SAXS data to bias simulations “on-the-fly”, as it was done in atomistic Molecular Dynamic simulations for proteins, to follow global rearrangements. When the structure of the molecule was available at full resolution in one state, SAXS data allowed to study an opening-closing transition involving the rearrangement of secondary structure elements [24, 25].

The situation however becomes immediately more challenging if the local structure is not unambiguously available for the whole molecule and one has to explore the conformational space widely, also in the quest of secondary structures. Because RNA molecules can adopt alternative conformations separated by high energetic barriers [26, 27], exploring the full set of 3D structures is extremely challenging computationally and is beyond the scope of most prediction methods. Coarse-grained models, simplifying the systems representation, have shown the potential for addressing this challenge [28, 29, 30, 31, 32, 33] and are therefore good candidates to be combined with SAXS data in this scenario. To avoid spending time to generate a full energy landscape to then focus on just one basin making a selection *a posteriori*, coarse-grained simulations can be combined with SAXS data in a strategy where experiments limit the search *a priori*, leading a simulation to explore only the region of the conformational space that is compatible with the data. For this purpose, the use of coarse-grained models coupled with SAXS experiments is particularly pertinent. In order to bias simulations towards structures compatible with SAXS data, multiple evaluations of the scattering curve are required throughout the trajectory. Using Debye theory [34], one needs to compute *N*^2^ terms, with *N* the number of particle of the system. A coarse-grained approach lowers significantly the number of particles and consequently reduces the computational costs, allowing the integration of the theoretical SAXS curve computation in the simulations more easily. We therefore exploit the coarse-grained representation to describe molecules with a lower number of effective beads with respect to the number of atoms in the atomistic representation, while keeping a level of resolution adequate for a comparison with SAXS data which is intrinsically at a low resolution.

We present here an approach based on coarse-grained modeling to make use of SAXS data in the context of the folding of RNA molecules, to help predicting systems for which parts of the local structure is unknown, with the experimental data guiding the folding of the molecule. This method takes advantage of the ability of our coarse-grained model HiRE-RNA to explore alternative structures and to fold from fully stretched configurations [33, 35, 36]. The SAXS bias is introduced as an additional energy term in the force field which guides the folding process to explore a particular region of the landscape while allowing the molecule to rearrange locally, following physical interactions and forming the base-pairing and stacking that ultimately define the folded structure even in the absence of the bias.

In section 2, we are going to present the theoretical framework used to build scattering intensity curves for a coarse-grained model and discuss the application to HiRE-RNA. We discuss in detail how going from an atomistic calculation of scattering curves to a coarse-grained calculation can induce errors that are dependent on the choice of the beads, and we propose a statistical correction for the coarse-grained calculation that recovers a full agreement with the atomistic curve. We then present in section 3 the strategy to integrate the comparison of a scattering curve computed on-the-fly in the simulation to a target curve. This consists in building a scoring function that is adequate to bias simulations aiming at obtaining folded structures from unfolded or partially unfolded initial conformations, which is different from a scoring function to perform the fit of a more rigid molecule, and on determining an efficient simulation approach. We find that the use of internal coordinates [37] to explore the energy landscape, combined with the smoothness provided by the coarse-grained model, is key to achieve folded structures from partially unfolded conformations. Finally, in section 4 we present the results of this approach on three benchmark systems for which we show that the addition of a biasing force based on the SAXS score significantly improves the efficiency of the simulation in finding structures with low RMSD and high base-pairing fidelity when compared to the known native structures.

## 2 SAXS intensities calculations

The formalism to obtain theoretical intensity SAXS curves for a biomolecule of a given structure is based on Debye scattering theory [34]. Despite the fact that this theory is rather straightforward, the details to make it applicable for a coarse-grained system are crucial and can be tricky at times and special care is required for a coarse-grained model. We therefore find it useful to briefly review this approach, starting with the implementation for an atomistic description of a system to then derive the theoretical framework for a CG representation, following the steps of what was done for proteins [38], and then present the special adjustments needed for our specific model.

### 2.1 Atomistic scattering intensities

The scattering amplitude *A* for a dilute sample receives a first contribution *A_v_* from the scattering amplitude *in vacuo* for the molecule, and a second contribution *A_s_* from the scattering amplitude of the solvent. It can also receive a third contribution *A_b_* from the scattering amplitude of the hydration shell. The detected scattering intensity can then be expressed as a function of the scattering momentum modulus 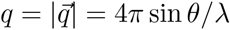 as

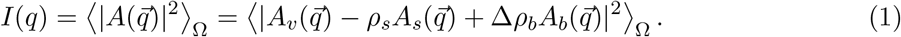

Here the brackets 〈...〉_Ω_ denotes the average over all orientations of the molecules, *ρ_s_* is the solvent electron density, and Δ*ρ_b_* is the contrast or excess density of the molecule with respect to the solvent.

Expressing the amplitude *A_v_* in terms of the atomic form factors *f_i_*’s, one obtains the Debye formula for the intensity *in vacuo* [34]:

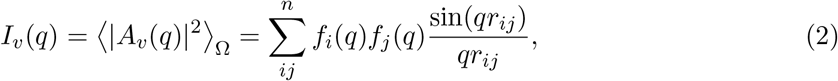

where the sum runs over the *n* atoms in the molecule, and *r_ij_* is the distance between atom *i* and *j*, 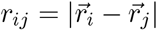. Adding the contribution from the excluded volume as in equation (1) modifies the effective form factors, yielding

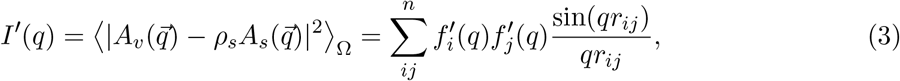

with, in good approximation [39],

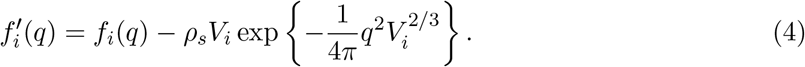

Here *V_i_* is the volume of the atom *i*, as observed by experiments (Table 1). To account for the hydration shell, an algorithm calculates a border layer of the molecule’s occupied volume and add its contribution to the intensity. The contrast parameter can be varied in order to find the best fit with the experimental curve, as it is done by CRYSOL [11].

**Table 1:**
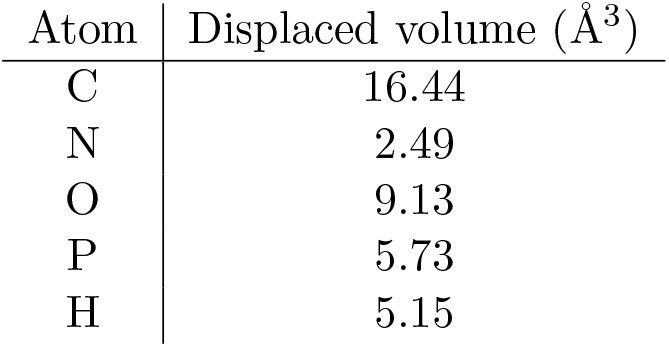
Excluded volume parameters for the atoms composing nucleotides [39, 11].

| Atom | Displaced volume ($\text{\AA}^3$) |
| --- | --- |
| C | 16.44 |
| N | 2.49 |
| O | 9.13 |
| P | 5.73 |
| H | 5.15 |

### 2.2 Coarse-grained scattering intensities

According to equations (1) and (3) to compute a scattering intensity curve from a coarse-grained structure, we first need to derive the expressions for form factors of the beads of the CG model. The intensity *in vacuo* for the coarse–grained molecules takes the form of the Debye formula:

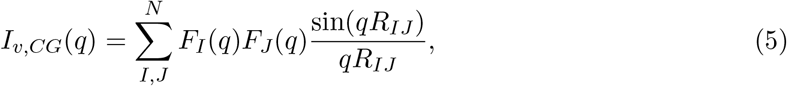

where the *F_I_*’s are the form factors for the molecule beads, and the sum runs over the *N* beads of the molecule, with *R_IJ_* the distance between the bead *I* and *J*, 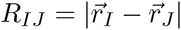. The coarse–grained form factors are calculated evaluating the scattering amplitude from the grain, yielding

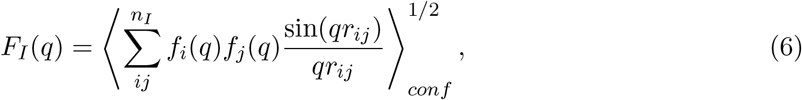

with the brackets 〈...〉_*conf*_ denoting the average over the relevant conformations of the atoms forming the bead. Here the sum is for *i*, *j* running over the indices of the *n_I_* atoms composing the bead *I*. There are no free parameters in the expression for the coarse–grain intensity in *vacuo I_CG_*.

The implicit solvent is taken into account by subtracting the excluded volume term from the atomic form factors, using equation (4), and substituting it in the formula for the modified coarse–grain form factors, in analogy to equation (6), yielding

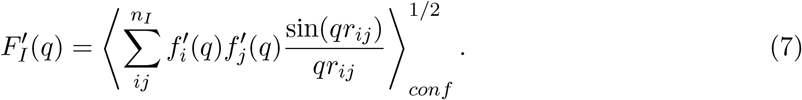

The equation above represents the subtraction of the solvent contribution from the scattering intensity.

Care is needed when taking the square root in equation (7). While the analytic expression given by equation (7) always gives positive values of coarse-grain form factors by definition, the sum determining the value at zero scattering vector,

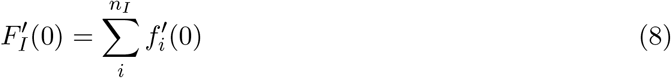

can yield negative values, due to the excluded volume subtraction. Should this be the case, one needs to consider the negative solution of the square root [40] and to correct the associate form factor in order to guarantee its physical meaning. In general, a polynomial interpolation is used to calculate the physical form factor, so that it is characterized by a monotonic growth for small *q*’s, and by the correct value at *q* = 0 [39].

In formulae, the *in vacuo* and implicit solvent terms read

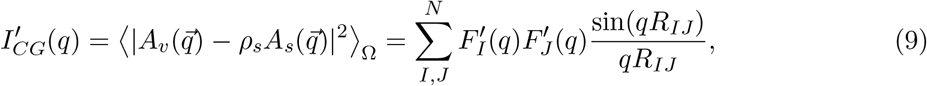

where the form factors are derived as explained above.

#### Hydration shell

We consider an explicit one-particle layer for the hydration shell, formed by water molecules located at the Oxygen atoms positions, with form factors 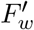, accounting for the buffer subtraction, given by

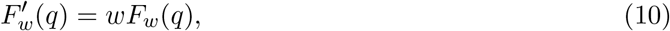

where *F_w_* is the tip3p [41] form factor for the water molecule. The parameter *w* is related to the excess electron density by

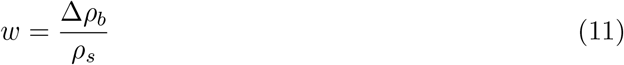

and can be varied in order to fit the experimental intensity curve. Typical values for *w* range from 0.01 to 0.3 [38, 11].

The total scattering intensity in solution for a coarse–grained molecule with *N* particles and *M* dummy water molecules in the hydration shell is then

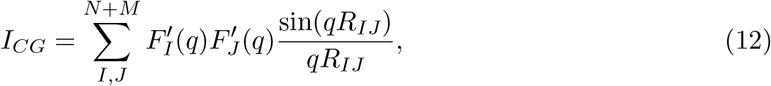

with 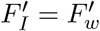 for *N* < *I* ≤ *N* + *M*.

### 2.3 Application to the HiRE-RNA coarse-grained model

In this section we are going to apply the results of coarse-grained scattering intensities calculations to the coarse-grained HiRE-RNA model that we develop. We then compare coarse–grain and atomistic results for a library of structures, in order to build a framework of applicability of our coarse-grained procedure.

HiRE-RNA is an implicit solvent, implicit ion model [42, 33] where each nucleotide is represented by 6 or 7 beads corresponding to the backbone heavy atoms P, O5’, C5’ and C4, C1’ of the sugar, and to the center of mass of each of the aromatic rings of the bases (G1, G2, A1, A2, C1, U1) (Figure 1). The model has been presented in all details in [33]. The force field of the model is composed of local interactions accounting for the local stereochemistry, an excluded volume interaction giving a physical size to the beads, and non-local interactions accounting for base pairings (short-range), base stacking (short range) and electrostatics (long-range). The model was developed with the specific aim of studying nucleic acids folding and assembly.

**Figure 1:**
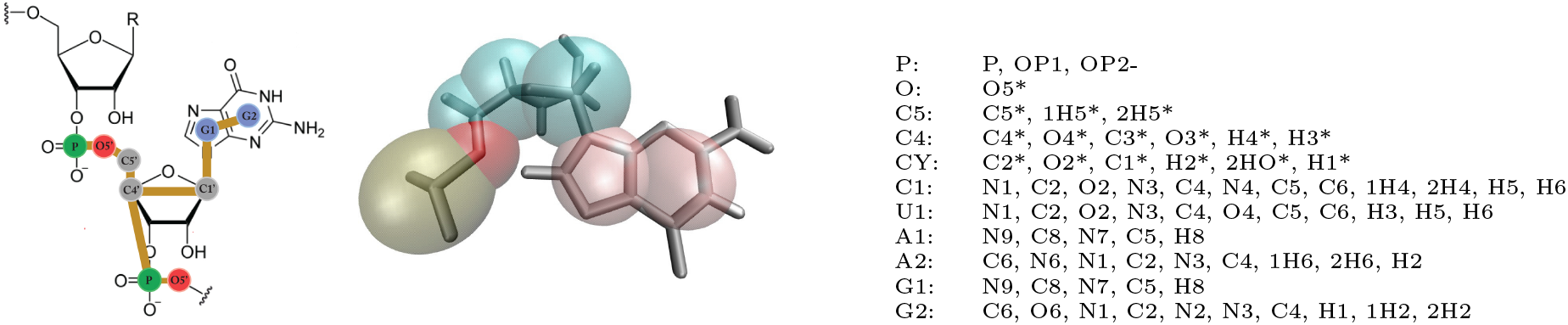
Definition of grains in HiRE-RNA model.

The *in–vacuo* and in-solution form factors for the grains of the HiRE-RNA model are calculated using formulæ(6) and (7) respectively. As mentioned in section 2.2, when buffer subtraction is considered, some form factors may need a more careful evaluation, in order to account for the correct root of the quadratic equation defining them. In our model this is the case for the bead C5 for which the derivation of the form factor is discussed in SI (appendix A.1).

To assess the validity of the coarse–grain intensity *in vacuo* and in solution, given by equations (5) and (12) respectively, we evaluate the difference between the coarse–grain and atomistic scattering curves for a library of 178 structures, characterized by different topologies and sizes. In both cases we find that there is a general trend in the discrepancy between the curve calculated from the atomistic description and for the CG description (figure 2, an example on a specific molecule is shown in SI, section A.2). The difference is small for small values of *q* increases for higher values of *q* showing a periodicity, and value of the logarithm of the intensity for the atomistic curve is always lower than that of the CG curve. To reduce the error introduced by coarse-graining, we compute the average difference

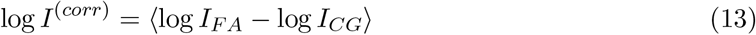

over all structures, and use it as a correction for the CG intensity defining:

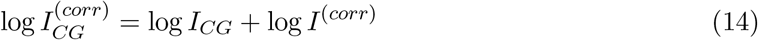

**Figure 2:**
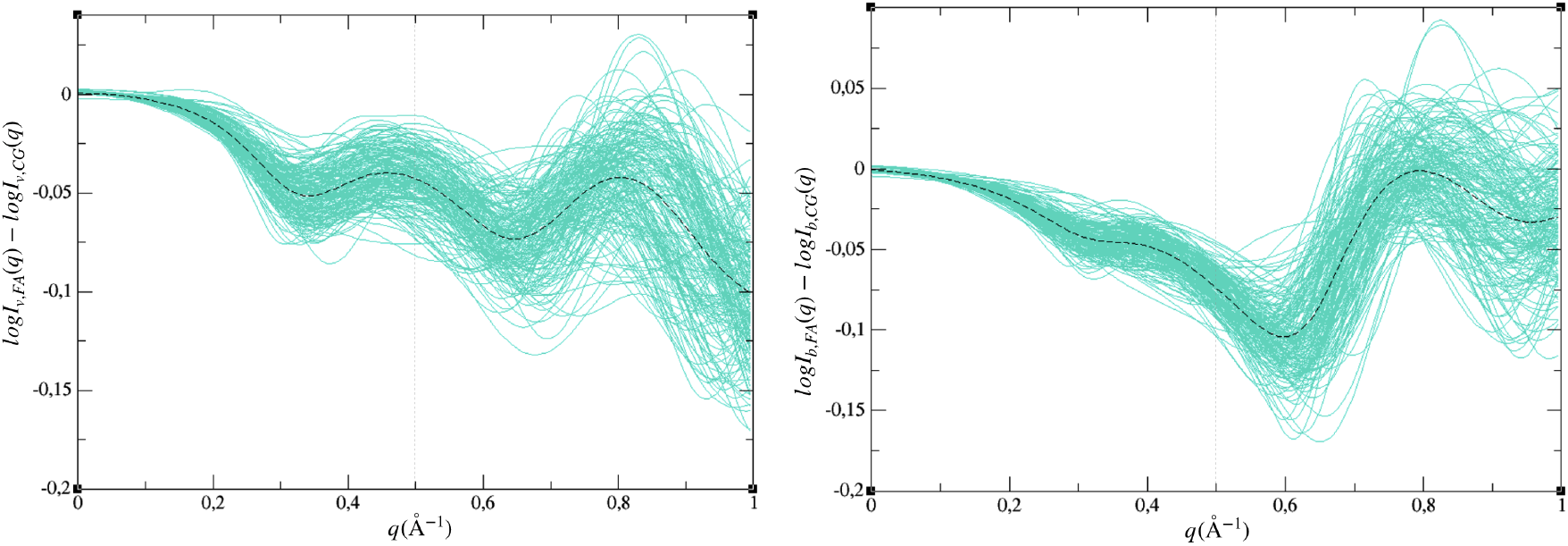
Difference between full–atom and coarse–grained intensities for 178 structures *in vacuo* (left) and with contrast (right). log *I*^(*corr*)^ is shown as dotted line.

#### Hydration shell

In order to compare our theoretical computation of the scattering intensity to experimental SAXS curves, we need to set a value for the excess electron density *w*, appearing in the equation for the water form factor of the hydration shell, equation (10). Comparing the theoretical curves computed over our library of structures with results from CRYSOL, we set *w* = 0.037, which yields the best agreement. The parameter defining the radius of the effective water grains is fixed to *r_w_* = 1.2*Å*, consistently with the size of the water molecule. Its value, in principle, could be varied, but it is correlated to the value of the shell contrast *w*.

The contribution of the hydration shell to the total scattering intensity is generally considered to be negligible [40, 43], when dealing with differences between states. In our scheme we will always be considering the difference between a target intensity curve and the istantaneous theoretical scattering curve computed during the simulation, using in particular the same mean correction *I*^(*corr*)^ defined in the buffer–subtracted framework, equation (14). In this case, in order to exploit the possibility to compute the hydration shell contribution to the intensity only once, this term should be evaluated for one conformation of the considered structure (namely the initial conformation for the simulation) and subtracted from the experimental curve, yielding an effective buffer–subtracted intensity. In formulæ, if *I_exp_* denotes the experimental intensity, *I_init_* the total intensity for the initial coarse–grained conformation for the simulation, and 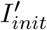 the buffer–subtracted intensity for the same conformation, the reference curve for buffer–subtracted calculation 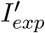 reads

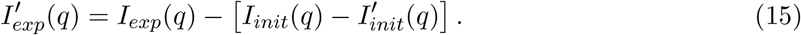

To verify that indeed we can avoid calculating the shell contribution during a simulation, we analyzed the trajectory of several MD simulations of four different molecules, for which we computed a score with the hydration shell updated at every step as the structure evolves and a score with the shell contribution taken always from the first frame of the simulation. We find a maximum difference between the two methods of at most 15% of the score value for RMSD deviations of more than 20 Å, which indicates that indeed we can conceive a strategy where we compute the hydration shell contributions only once (an example is shown in SI, section A.3). This implies a big gain in terms of computational costs, since the algorithm needed to compute the shell and its contribution to the scattering intensity is quite expensive, adding *M*^2^ terms, with *M* > *N*, with *M* the number of solvent grains and *N* the number of grains in the molecule.

## 3 Guiding coarse–grained simulations

In order to guide simulations towards the target structure by means of low–resolution data, we define a score based on the difference between the computed and the target intensities. When multiplied by a constant to make its numerical value comparable with the internal energies of the system, we interpret this score as a “SAXS potential energy”, *E_SAXS_*, from which we compute a “SAXS force”, *F_SAXS_*. This biasing potential is evaluated throughout the simulation and the force can be added as a bias in minimizations and in MD.

### 3.1 SAXS energy and SAXS force

Several scoring functions have been proposed in the past [11, 44, 45, 46] to compare SAXS intensities curves. Having tested all of these functions with our simulation scheme, we found that none of them was suited to capture the difference in intensity curves for the comparison between a partially folded RNA and its native structure. This is because the weight of the score was either too skewed toward very low values of *q* or toward high values of *q*, while values of *q* between 0.2 and 0.4 Å^−1^ plaid little role. However this is the region were we observe the largest differences in the curves for a partially unfolded structure. We therefore defined the following score, which gives sensibly better results when comparing the coarse–grained computation to a target curve:

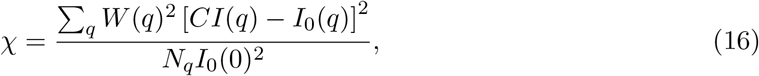

with

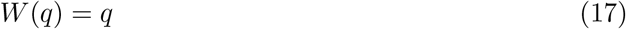

In the above expressions, equation (16), *I*_0_(*q*) denotes the target intensity curve, while *I*(*q*) is the intensity calculated during the simulation, as a function of the scattering momentum *q*. Since we are interested in fitting to low–resolution data, the sum limits for *q* will be *q*_0_ = 0 and *q*_1_ = 0.5 Å^−1^, while *N_q_* is the number of points for which the intensity is evaluated. The linear weighting function, equation (17), provides a larger weight for intermediate values for the scattering vector, which is what we seek. The constant *C* allows the comparison with experimental data and is related to the rescaling of the scattering intensity due to the sample dilution. In general, the rescaling of the simulation scattering intensity amounts to

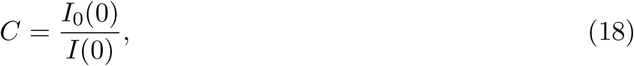

ensuring the same scale for the experimental target curve and the theoretically calculated intensity^1^.

From equation (16) we define the SAXS biasing potential as

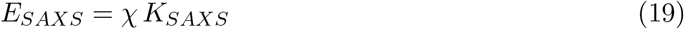

Where *K_SAXS_* is a constant carrying the dimentions of an energy. With our definitions, beacuse of the normalization of *χ* by *N_q_*, typical values of *K_SAXS_* are of the order of 10^6^ - 10^8^ to make *E_SAXS_* of the order of the unity and make it comparable with the energies of the force field.

The SAXS force, actively drawing the simulated molecule’s SAXS curve to fit the target scattering intensity, can be computed as the gradient of the SAXS energy over the relevant spatial variables, which in this case are the coordinates of all the particles in the system. It is therefore a force acting at once on the whole system.

From equation (19) we calculate the force acting on the particle *I* as

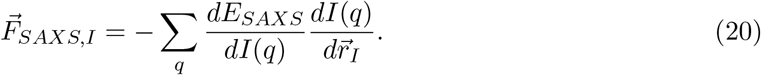

Using the definition of the SAXS intensity, equation (12), the two terms in the expressions above are given by:

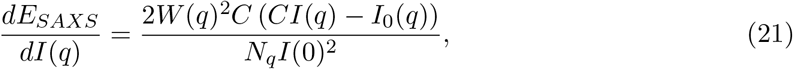

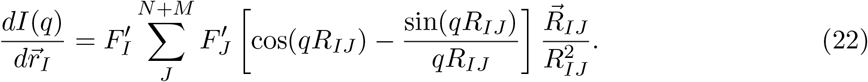

Substituting our definitions for the weighting function *W*(*q*) and the constant *C*, equation (17), we arrive at the SAXS force for a theoretical target intensity.

### 3.2 Biasing strategy

Our goal is to guide nucleic acids simulations towards the corresponding native structure, exploiting the knowledge of the associate SAXS profile, starting from partially or completely unfolded conformations. We present here what we found to be the most efficient strategy among the many we tested.

At first we have to scale the *E_SAXS_* to values that are comparable with the energies of the force-field in order for the bias to be of some effect. If, for the initial configuration, *E_SAXS_* is too small the biasing effect is not enough to drive the simulation, while if it is too high the biasing effect dominates over the force-field and the internal structures of the molecule are deformed. Starting from a relaxed initial conformation (usually partially unfolded), we perform an energy minimization using quasi-Newton optimization [47] with internal coordinates [37] including *E_SAXS_*. This is followed by an additional unbiased relaxation though a quick coarse-grained molecular dynamic simulation. For sample frames in the MD trajectory we compute *E_SAXS_ a posteriori* and the number of paired bases and select the structure with the lowest score and the highest number of base-pairs (since we want to go from a partially unfolded to a fully folded structure). We then start an iterative process in which the selected frame is again minimized with the SAXS bias followed by an unbiased relaxation MD. We stop the process when we observe no further gain in lowering *E_SAXS_* and in adding new base-pairs. This typically occurs after 2 or 3 cycles. The best final conformation according to these criteria is then converted from coarse-grained to all-atom representation in order to evaluate its stability through a longer (several hundreds ns) molecular dynamic simulation without constraints. The resulting trajectory is then clusterized to compare the most representative conformation with the reference structure. Figure 3 illustrate the whole approach.

**Figure 3:**
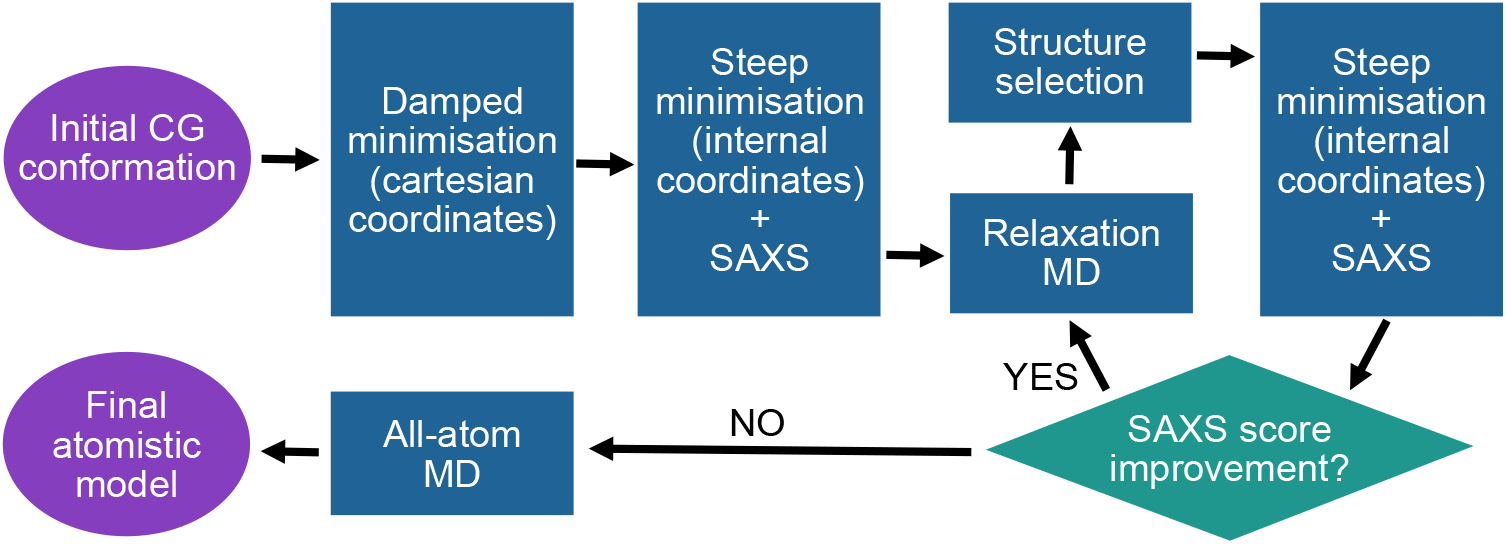
Workflow for the iterated rounds of SAXS–guided minimizations, alternated with relaxation MD’s: from the initial conformation to the final candidate structures compatible with SAXS data.

We find that the use of internal coordinates in the minimization is crucial because it allows the structure to explore largely diverse conformations without distorting the molecule by stretching bonds and angles, which would be very costly energetically. It therefore ensures a smoother minimization process able to reach minima with greater structural differences from the initial structure than a minimisation in Cartesian coordinates [37]. The details of the implementation of internal coordinate minimization to HiRE-RNA are presented in SI (appendix B).

The value of *K_SAXS_*, and therefore the scale of biasing potential, can be adjusted from one iteration to the next so that *E_SAXS_* remains of the same order of magnitude as the energies of the force-field, and can also vary from one system to another depending on the stability of the initial structure.

To assess the gain given by the fitting procedure to SAXS data, as opposed to the simple use of the CG force-field for folding, for all systems we perform a control 150 ns MD simulation in the CG representation, to allow the system to reach the native state without bias.

## 4 Results on benchmark systems

In this work we want to show that in principle our approach can lead to native-like folded structures from partially unfolded conformations. We have chosen 3 benchmark molecules for which we have built target scattering intensity profiles theoretically from the known native high-resolution structure and for which we generated partially unfolded structures.

The benchmark molecules range in size from 44 to 77 nucleotides and include 2 pseudoknots. Their main features are presented in table 2. The initial conformations for our simulations are partially unfolded structures. The idea is that using other experimental information or other modeling tools such as secondary structure predictions, we can often determine some of the base pairings and local structures of the molecule, but not the fully folded structure. For the two largest molecules, partially unfolded structures were obtained by manual unfolding from their native state using interactive simulations performed with the software UnityMol [48]. By pulling on different beads, local structures were broken and various parts driven apart from each other. For the smallest molecule we applied what would be the typical procedure of determining an RNA structure from the sequence alone: using [14] we predict the secondary structure and using RNA Composer [15] we obtain a 3D structure. Depending on the predicted base pairings, this structure might be only partially folded.

**Table 2:** Summary of the main properties of our benchmark molecules: PDB ID, complex structure and function, experimental method (Exp.), total number of nucleotides (Nt), total number of base pairs (BP), kind of secondary structure, structural diffrence of the partially unfolded state.

| *PDB ID* | *Structure* | *Exp.* | *Nt* | *BP<sup>a</sup>* | *2D structure* | *RMSD<sup>b</sup>* |
| --- | --- | --- | --- | --- | --- | --- |
| **1A60** | Portion of tRNA | NMR | 44 | 16 | H-Pseudoknot | 15.8Å |
| **1P5O** | Domain II of HCV IRES | NMR | 77 | 32 | Stem-Helix | 16.0Å |
| **1Y26** | Riboswitch | X-Ray | 71 | 30 | kissing loops | 22.4Å |

<sup>a</sup> Base-pairings are referred to x3dna webserver including 1Hbond; <sup>b</sup> of the partially unfolded initial structure

For each system, the target SAXS curve is calculated using the same approach of sections 2, which leads to intensity curves in excellent agreement to those produced by the program CRYSOL [11]^2^. Since the contribution from the hydration shell to the total scattering intensity can be computed only once, rather than at each step of the imposed SAXS bias, we evaluate the scattering curve for the target curve corresponding to the buffer–subtracted intensity given by equation (9), with the mean correction introduced in equation (14). Consequently, the evaluations throughout the simulations are also performed in the buffer–subtracted framework, rather than with the addition of the hydration shell, allowing to save a sensible amount of time and computational resources.

The biasing strategy presented in section 3.2 is applied to the 3 benchmark molecules leading to the scattering curves of figure 4. We detail below the structural content for each molecule.

**Figure 4:**
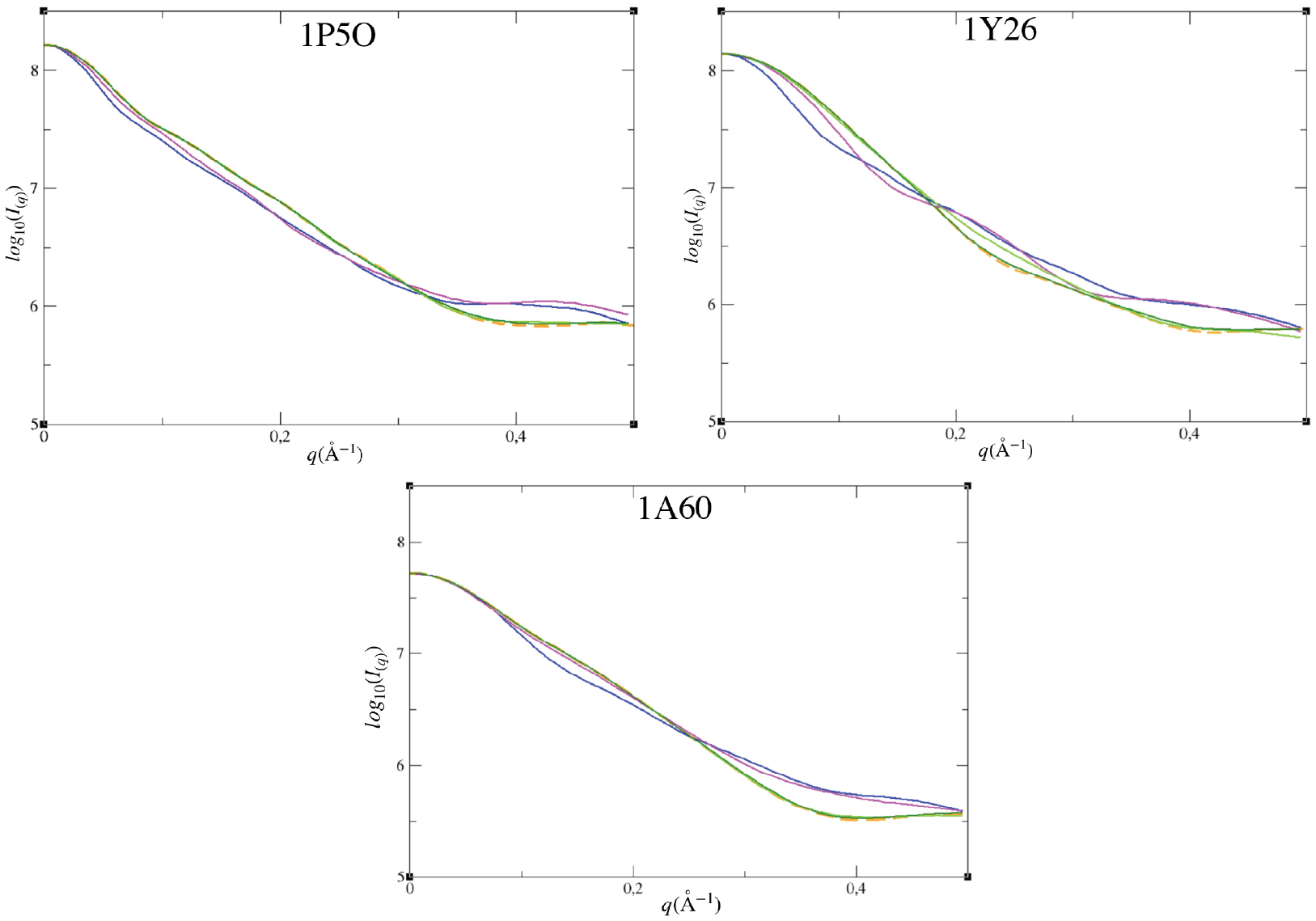
Computed scattering intensity curves for the three structures: native structure (orange), initial partially unfolded structure (blue), control structure from unbiased MD (purple), resulting structure from the first biasing minimization (light green), structure resulting from all cycles of SAXS biasing coarse-grained modeling (dark green)

### 1P5O

The partially unfolded conformation of 1P5O has 20 base pairs, 19 of which in common with the 32 base pairs of the native structure, and has an RMSD of 16 Å. A *K_SAXS_* equals to 10^8^ gives an initial *E_SAXS_* of about 600% of the internal energy at the beginning of the first minimization, which reduces to 0.5% at the end of the minimization. At the end of the relaxation MD performed after the first minimization (end of the first cycle), the number base pairs increases to 27. In Figure 4:1P5O we can observe the progression of the intensity curves and we see that the curve after the first iteration is closer to the that of the target, but there is a visible discrepancy for values of *q* greater than 0.2 Å^−1^. We therefore continue with another cycle of biasing using the same values of *K_SAXS_*. At the beginning of the second minimization *E_SAXS_* corresponds to 75% of the internal energy and reduces to 0.6% at the end of the second minimization. At the end of the two cycles the score *χ* has reduced from 7 × 10^−6^ to 3 × 10^−9^ and the intensity curve of our minimized structure is almost indistiguishable from the target in the full range of *q*. The number of base pairs increases further to 31, 28 of which in common with the native structure. Three base-pairs are slightly misplaced by one base shift in our final structure (see SI D1: 1P5O Table 3). The RMSD from the native structure at this stage is of 7.8 Å.

The structure issued from the second minimization/relaxation cycle is then converted from coarse-grained to all-atom and the MD is performed. The middle structure of the most representative cluster during all-atom trajectory has a RMSD of 5.6 Å and contains 32 base pairs, 30 of which in common with the native structure and 2 of them shifted by one base. It is worth mentioning, that 9 of the native base pairs are non-canonical and that our procedure is able to correctly recover 7 of them. The final structure is shown in figure 5:1P5O-C. The evolution of base pairing at the different cycles is shown in SI D1: Table 1P5O. For comparison, the final structure of the control simulation without SAXS bias has a significantly different shape from the native structure and from the structure obtained with our biasing procedure (figure 5:1P5O-B). It’s RMSD is of 11.4 Å, it has 30 base-pairs, 27 corresponding to native and 3 off by one or two bases.

**Figure 5:**
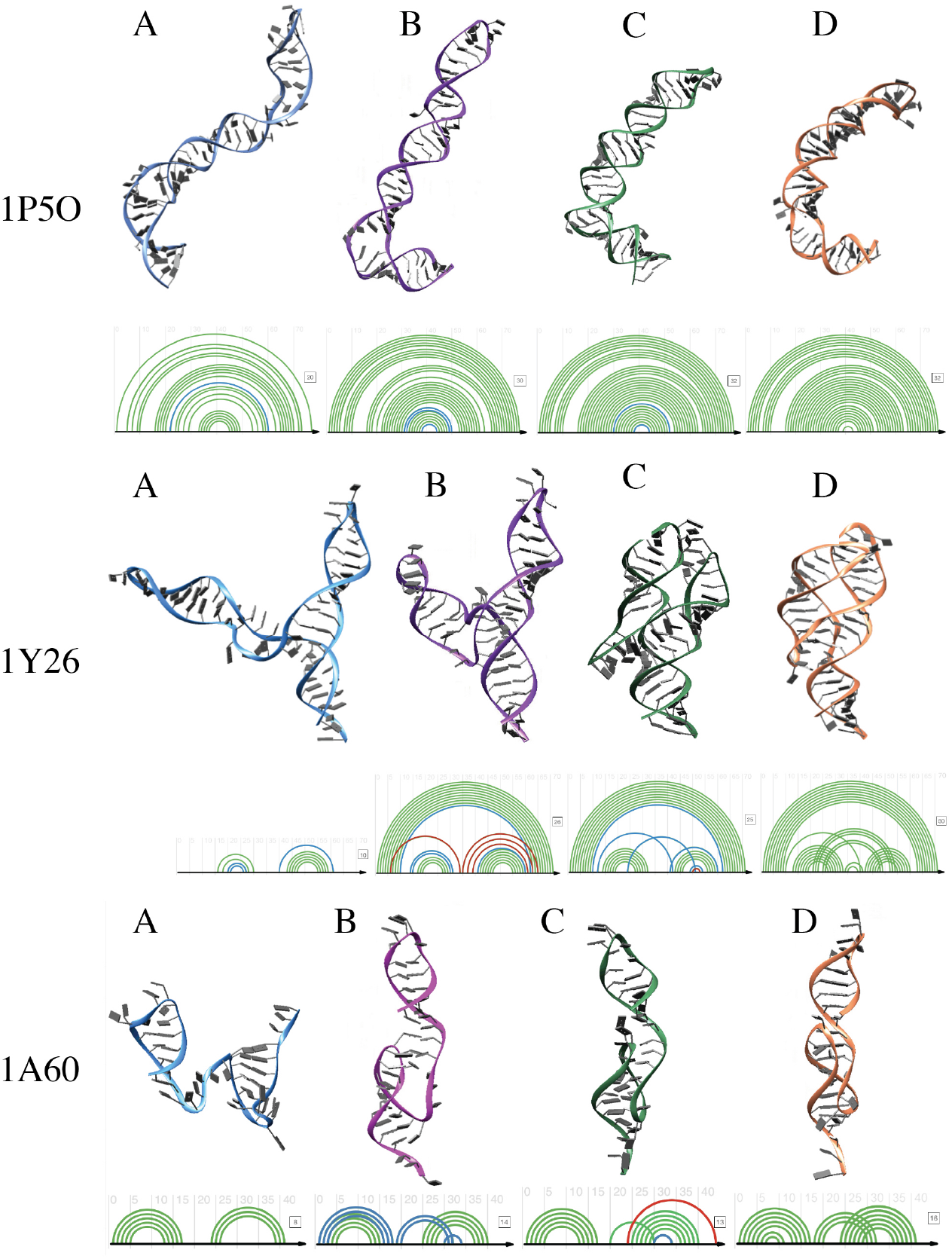
3D conformations and base paring arc diagrams [49] for the three molecules: A) partially unfolded initial structure, B) control structure, C) retained structure from SAXS-biasing, D) native structure. For each structure arc diagrams show native base pairs (green), base pairs shifted from one or two bases from native (blue), and base pairs shifted from the native beyond two bases (red).

### 1Y26

The partially unfolded conformation of 1Y26 contains 10 base pairs, 7 of which in common with the native structure, and has an initial RMSD from native of 22.4 Å. Given the relative small portion of paired bases out of the full length of the molecule (20 out of 77), the structure is not very stable and we begin the minimization procedure with a softer bias compared to the previous case, setting *K_SAXS_* equals to 10^6^. This gives a biasing potential *E_SAXS_* of 50% of the internal energy. At the end of the first minimization, *E_SAXS_* reduces to 1% of the internal energy, with a *χ* score of 6.6 × 10^−7^. The number of base-pairs doubles, with 11 of them in common with the native structure and 9 not corresponding to native. The intensity curve overlaps with the target for *q* < 0.2 Å^−1^, but departs for higher values.

For the second cycle we increase the biasing, setting *K_SAXS_* = 10^8^, which gives *E_SAXS_* of 570% of the internal energy. At the end of the second minimization *E_SAXS_* reduces to 1% of the internal energy and *χ* = 9 × 10^−9^. 23 base pairs are formed, 19 of which in common with the native structure. The two arms of the structure, that in the native configuration form a kissing loop, are now close to each other and one base pair is formed across the loops between bases 24 and 51. The intensity curve at this stage overlaps the target for the full range of *q* (figure 4:1Y26).

The resulting conformation is converted from coarse-grained to all-atom and a 300ns MD trajectory is analyzed. The most populated cluster of the dynamics has an RMSD of 6.9Å and forms 25 base pairs, 19 of which in common with the native structure, 5 off by one or two bases, and 1 off by more than two bases. One of the 5 non-canonical base pairs is recovered. The evolution of base pairing at the different cycles is shown in SI D2: Table 1Y26. The overall architecture of the molecule is recovered and one base pair hold together the kissing loop. It is to be noticed that the native experimental structure for this molecule was obtained with the ligand (adenosine) in the central junction, while in our simulations the ligand is missing and therefore we expect some local variations from the native structure.

In the control simulations the kissing loop is never formed and the two inner stems remain far apart. The external stem forms spontaneously and non-native base paris elongate the two inner stems. The final RMSD is of 15.8Å and the number of base pairs is 26, 18 of which native, 4 off by one or two bases and 4 off by more than two bases.

### 1A60

The partially folded conformation of 1A60, obtained from RNAfold and RNA Composer from the sequence as only input, presents two paired regions forming stems/loops, connected by an unpaired strand. 8 base pairs are present, all in common with the native structure, which has 16 base pairs overall. Because of the lack of structure of the initial configuration in the central, unpaired, region, and of our procedure that strongly depends on a minimization from the initial configuration, we found that the success of the fit were dependent on the choice of initial structure. We therefore decided to generate three alternative structures from the one proposed by RNA Composer, applying random torsions to the unstructured central strand. The three structures, named S1, S2 and S3, have RMSDs of 18.9Å, 20.4Å and 15.4Å respectively and initial *χ* values of 1.2 × 10^−5^ for S1 and 9.6 × 10^−6^ for S2 and S3. For each of them we apply our procedure and compare the results at each step.

After the first minimization S1 formed 15 base pairs and *χ* = 2.3 × 10^−8^, S2 formed 12 base pairs and *χ* = 7.3 × 10^−9^, and S3 formed 13 base pairs and *χ* = 5.5 × 10^−9^. By visual inspection we observe that the fold achieved by S1 does not resemble the typical structure of an RNA, with important deformations of the local structures, including the hairpins. Moreover, because its fit to SAXS data is worse than for S2 and S3, we retain only S2 and S3 as possible candidates that we submit to the second cycle. At the end of the second relaxation, S2 formed 16 base pairs and *χ* = 4.0 × 10^−9^ and S3 formed 18 base pairs and *χ* = 8.3 × 10^−9^. At this stage S2 also exhibits deformed double stranded regions and it is less stable than S3, with an internal energy twice as high. This process bring us to retain only S3 for the final atomistic MD.

The final structure has and RMSD of 6.3 Å and 13 base pairs 11 of which in common with the native structure, one with one or two bases difference, and one with more than two bases difference. This last base pair occurs as the dangling end, which is unpaired in the native structure, interacts with a base near by in our case. The overall architecture of the molecule is recovered with the pseudoknot forming correctly, although stabilized by fewer base-pairs.

The control simulation with no bias also achieved a folded structure with the correct pseudoknot (RMSD of 7.0 Å, 14 base pairs, 8 native and 6 off by one base), highlighting how for relatively small molecules the force field alone is able to obtain the correct structure, however the time needed for this results is much higher as the fold is reached after 50 ns of simulation, requiring two days of computation.

## 5 Discussion

We have shown here that it is possible to develop a theoretical framework to include a SAXS bias onto coarse-grained RNA simulation implementing a successful strategy for structure prediction. This is achieved using Debye theory for scattering adapted to the coarse-grained model by the calculation of modified form factors, the inclusion of a statistically computed correction term, which becomes important at intermediate and high values of scattered momentum *q* where the difference between the atomistic and the coarse-grained representation is significant, and by the evaluation of solvation by a coarse grained approach on the water as well, which can be computed once and for all for the molecule and applied without modification throughout the simulation, therefore speeding up computations significantly. The comparison between SAXS profiles evaluated on-the-fly in the simulation and a target intensity curve is done via a scoring function chosen to give a significant weight to the portion of the curves that is relevant in the coarse-grained representation, that is up to values of *q* of about 0.4 Å^−1^. This is also the limit after which typically the experimental curves becomes too noisy to be used. This score is then interpreted as an energy and it is possible to compute analytically a force associated to this term that can be added to the force-field. As opposed to the forces of the particles interactions, which act at the most on 6 particles at the time in our model, this force is global, in the sense that it concerns all particles at once. Its effect can be imposed on the system via a coupling constant regulating its strength with respect to the force field alone.

We have designed a strategy to apply this force as a bias in our simulations using cycles of minimizations in internal coordinates and short coarse-grained MD to relax the system. This is the result of several prior attempts to design a successful strategy. The first unsuccessful attempt was to run MD simulations with the SAXS force active in the production phase either at all times or at fixed intervals once the structures has significantly changed from a previous evaluation. However, because of the time required to compute this term, even in the coarsegrained representation, the first option was too expensive and simulations too slow to converge to anything and in the second, evaluation at fixed intervals gave a contribution too weak to drive the simulation. A good balance between the two was not found. As a second attempt was to use a SAXS bias in a Replica Exchange simulation, where the SAXS energy is added to the internal energy in the Metropolis criterion. This option lead to a vague bias toward the native structure after several hundred ns of simulation, but failed to give satisfying results and was abandoned also because of its very high computational cost.

The scheme we have chosen, and presented here, takes advantage of the minimization in internal coordinates, which, while remaining basically a steepest-gradient descent, is able to modify the initial structure significantly thanks to the fact that only torsions are affected, while bonds and angles remain unchanged. This smooths out the landscape and allows much wider explorations than what it is possible in a minimization in Cartesian coordinates. The use of internal coordinates was therefore key to successfully introduce the SAXS bias. These minimizations have the advantage of being very fast, taking less than half an hour on a single CPU, and are therefore very affordable. Already after one such minimization we obtain structures that resemble the target structure, having adopted the overall shape. The following MD and the second minimization allow the system to rearrange locally, optimizing the internal interactions and forming the contact that ultimately hold the molecule together also in the absence of the bias, mainly by the formation of base pairs and tertiary contacts. A final atomistic MD allows the refinement of the structure and the possibility to verify the stability of the structure obtained after the minimization cycles.

For small molecules, secondary structure predictions and 3D structure predictions based on it are expected to give reliable results and are the standard used in RNA prediction. For our smallest molecule we therefore built the initial configuration with a procedure reflecting what is commonly done in the community. We show that even when the starting structure proposed by these methods is far from the native structure and is still partially unfolded, our implementation of the SAXS bias is able to recover the overall architecture and the key base pairs that hold the structure together. In some cases this might imply generating a few different initial structures to help the minimization reach alternative conformations.

We also compared our results with what could be the typical procedure using easily accessible and widespread tools, that is generating an envelope using the programs provided in ATSAS [50] and performing a fit into the envelope with normal modes, using for example the program iMODFIT [51] that is easily integrated in the visualization software Chimera [52] (work presented in SI section E). The first observation is that the straightforward application of the fit from a partially unfolded structure does not give satisfying results. The situation improves when one allows the system to form local structures by a short atomistic MD simulation (10 ns), followed by another fit to the envelope. Results obtained with this method are poorer in quality than what we achieve, lacking the base pairing details and therefore obtaining structures that do not hold together as the SAXS constraint is released. Moreover, this method is definitely more expensive than our approach, with results requiring the use of high performance computational facilities to perform the atomistic MD.

Overall, our results are still improvable, but are comparable in quality with the predictions of RNA puzzle competitions [17], with RMSDs a few Å from the experimental structure and a recovery of the correct secondary structure network of the order of 70-80%. It is important to mention that our predictions include also non-canonical pairs, which is often the weak spot of RNA 3D structure predictions, and that they capture the overall topology of the molecules, which is an important step for any further refinement. We believe the remaining discrepancy with the native structure is mainly due to the force field that still requires optimization (which is ongoing), implying that the implementation of the SAXS bias itself is promising.

As a next step, we will apply the same theory to bias Basin hopping Monte Carlo simulations [53] with our coarse-grained model. This will lead us to investigate structures more widely than what we can do in the current setup and it should to overcome the need of generating multiple initial conditions and make selections along the way. A large exploration of the energy landscape will also naturally include the possibility to consider the cases where multiple conformers of the molecule are present simultaneously in the sample, as we would have access to the different structures as the simulation evolves, allowing envision biasing strategies accounting for the visited structures with evolving relative weights and appropriate deconvolutions of the experimental signal.

As final remark, the theory and the practical approach that we presented here is for all purposes independent of the force-field used and, having worked out all the details of the implementation, can be easily extended to other models, for both RNA or proteins. Only the coupling between force field and SAXS contributions would have to be adjusted to satisfy the energetic balance between the internal energy and the bias. We consider this work as a proof of principle that on-the-fly SAXS biasing to generate RNA folds is a viable route and the approach we propose here could be included in integrative modeling programs considering multiple data sources and models at various resolutions.

## Acknowledgments

We are grateful for the support from the French National Computing facilities through grant A0090710584. L.A. thanks the doctoral school MTCI of the University of Paris for financial support. S.P. thanks F. Nitti for useful discussions on the theoretical development.

## Footnotes

1 For a comparison with experimental data with given errors *σ*(*q*), the choice of the scoring function should also include experimental errors in the denominator of *χ*, as done by the score defined in CRYSOL : 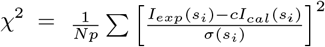.

2 Minor differences arise at very small values of *q* because CRYSOL attributes a neutral charge to the phosphate group of nucleotides instead of the correct negative charge at one of the oxygens.

## Notes

### Competing Interest Statement

The authors have declared no competing interest.

