## Supplementary material for "Biasing RNA coarse-grained folding simulations with Small–Angle X–ray Scattering (SAXS) data"

March 28, 2021

### A Theory details and examples

#### A.1 C5 form factor

In our CG model, C5 is a small bead containing 1 carbon atom and 2 hydrogen atoms and its form factor yields a negative value at zero scattering vector, as computed by summing the electronic densities of the corresponding atoms, $F_{C_5}(0) = -0,9322$. The physical part of the coarse–grain form factor, which is defined by the values of the scattering vector larger that the point where the form factor exhibits a minimum at low $q$'s, together with the $q = 0$ value, is fitted with a sixth order polynomial curve. The fit is performed considering the following three relevant points in the form factor curve:

1. the value of the form factor at $q = 0$;
2. a point close to the minimum of the curve at low $q$'s;
3. the maximum of the curve.

We use the resulting polynomial curve, shown in figure 1, is then used as the correct definition of the coarse-grain form factor.

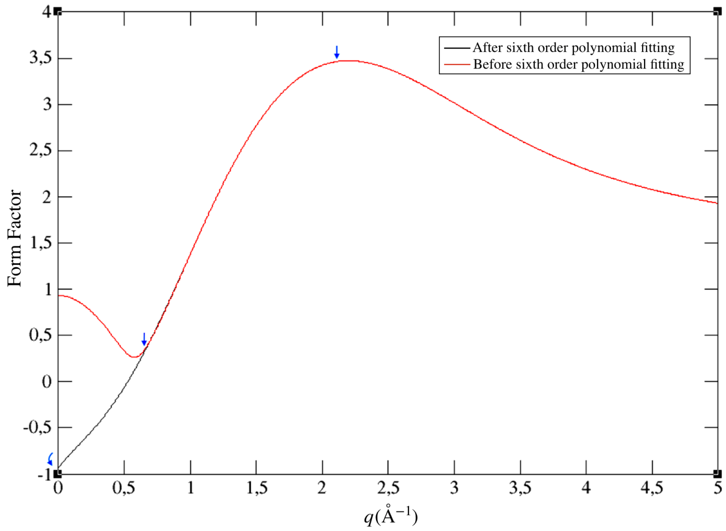

Figure 1: Corrected form factor of C5 grain obtained by interpolation with a sixth order polynomial curve. Blue arrows correspond to three points used for generating new curve equation.

The remaining grains do not exhibit this property and the corresponding buffer–subtracted form factors are directly given by the definition expressed by equation (7) of the main text.

#### A.2 Comparison of theoretical SAXS curves

In Figure 2 we report the SAXS intensity curve $I(q)$ for a specific molecule (1CQL, 43 nucleotides) as an example of the discrepancy in the calculations when using the atomistic model, the coarse-grained model without any correction, and the coarse grained model corrected with the curve obtained from the analysis of 178 structures. The difference in the curves becomes significant at values of $q > 0.2\,\mathring{A}^{-1}$. The quantitative difference between the curves in terms of the score $\chi$ defined in section 3.1 is $4.0 \times 10^{-7}$ for the uncorrected coarse-grained curve and $6.5 \times 10^{-8}$ for the corrected curve.

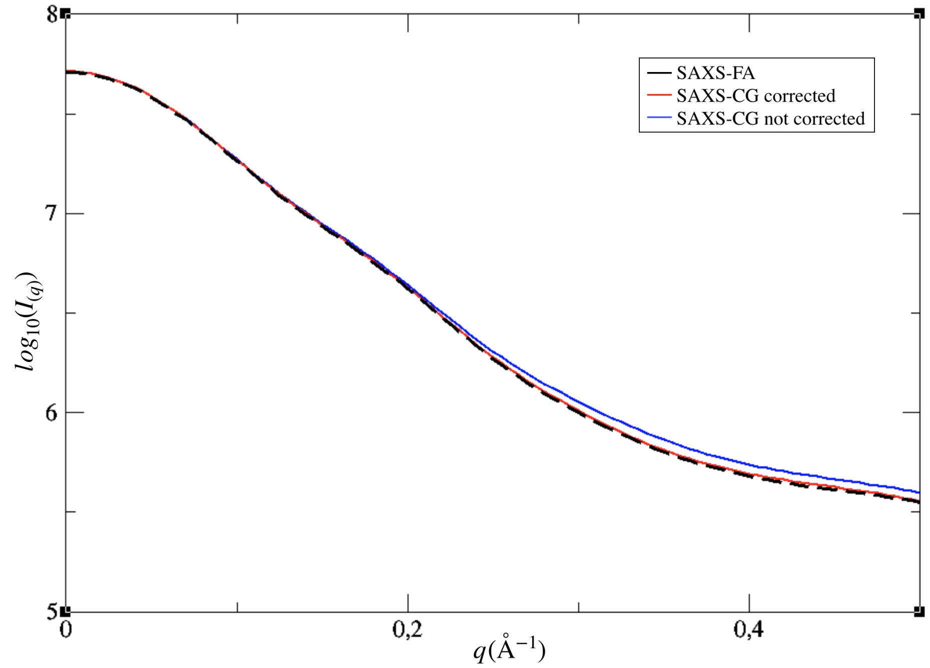

Figure 2: SAXS intensity curves for PDB ID 1CQL computed in vacuo with the atomistic model using Debye theory (black) and computed with HiRE-RNA coarse–grained model without any correction (blue) and corrected (red) using equation (14) (see main text).

#### A.3 Hydration Shell contribution

In Figure 3 we report an example of the effect of computing the shell contribution at every SAXS evaluation along a trajectory compared to the effect of computing it only once for the initial configuration, using this same contribution for all frames. The aim is to assess if the shell contribution changes significantly as the molecule explores different conformations. For this purpose we ran a long, unbiased, coarse-grained MD and computed the SAXS score $\chi$ *a posteriori* on a selection of frames with respect to the target curve obtained from the native structure. In making this comparison the normalizations in the definition of $\chi$ must be handled with care as for the simulation in which the shell is evaluated at every step the number of water molecules varies from one configuration to another. We therefore set $C = \frac{I_0(0)}{I(0)}$ to be able to capture the difference in shape of the two curves and not their overall size. The plot shows this score vs. the RMSD computed with respect to the initial configuration. As shown by the high RMSD values the molecules explore widely different structures, but the difference in $\chi$ computed with the two methods remains always of a few percentages of its total value.

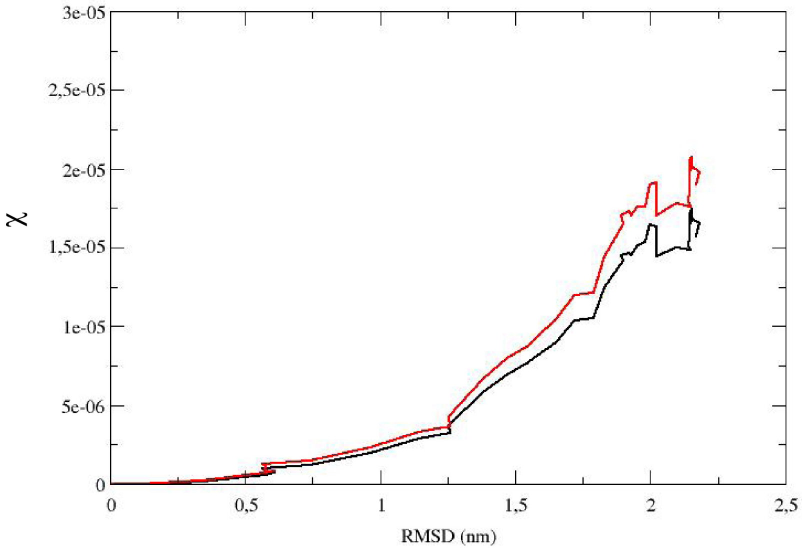

Figure 3: Variation of $\chi$ according to the RMSD from the initial structure for a coarse-grained trajectory of 1Y26 with the evaluation of the hydration shell at each frame (black) and at the initial frame only (red).

### B Internal coordinate minimization

We developed an internal coordinate minimizer using HiRE-RNA force field. To avoid artefacts, we chose the P atom closest to the centre of mass as pivot atom. The pivot atom is kept fixed during the minimization. We separated the backbone and base dihedral angles in order to be able to choose to minimize a given subset of dihedral angles ($\tau_i$). We read the atoms in the direction $5'$ to $3'$. For backbone dihedrals ($\tau_b$), if the rotation is made around a bond after the pivot atom, only the atoms after the bond are moved (rotation direction 3′ in Figure 4). If the rotation is made around a bond before the pivot atom, the rotation is characterized by the inverse order (rotation direction $5'$ in Figure 4), hence only the atoms before the bond are moved. For base dihedrals ($\tau_s$), the rotation direction is from the sugar to the last atom of the nucleotide base.

The derivative of the energy with respect to the independent variables $\tau_i$ and $\theta_i$ can be written as:

$$\frac{dE}{d\tau_i} = \sum_k \frac{dE_k}{d\mathbf{R}_k} \cdot \frac{d\mathbf{R}_k}{d\tau_i} \tag{1}$$

where the sum runs over the subset of atoms moving when a rotation is made around the unit bond vector $\mathbf{b}_i = (\mathbf{R}_{i+1} - \mathbf{R}_i)/s_i$, with $s_i = |\mathbf{R}_{i+1} - \mathbf{R}_i|$ for the torsional angle. This approach uses a modified Newton minimizer (Harwell VA13A) [1].

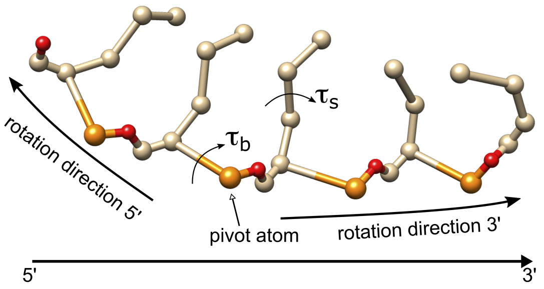

Figure 4: Definition of the sense of the rotation for backbone dihedrals.

To compute internal coordinate minimization, we use a using a quasi-Newton optimization algorithm (Harwell VA13A) which requires analytical derivatives of the conformational energy with respect to all independent variables, $x = x_1, x_2, .., x_n$. This algorithm uses the Broyden-Fletcher-Goldfarb-Shanno (BFGS) variable metric method without exact line searches of the type analyzed by Powell [1]. BFGS determines the descent direction by preconditioning the gradient with curvature information. In this method, the Hessian matrix is computed from the first derivatives and is gradually improved along the minimization. Convergence can be checked by observing the norm of the gradient, $\|\nabla f(x_k)\|$. Since the updates of the BFGS curvature matrix do not require matrix inversion, its computational complexity is only $O(n^2)$, compared to $O(n^3)$ in Newton's method. In the algorithm, two parameters have to be set: the scale (SCALE) and the accuracy (ACC). SCALE is a real array of at least $n$ elements, whose i-th component must be set to a positive value that modifies suitably $x_i$ initially used in the minimization calculation. About 10% of the total expected change in $x_i$ is often a good value. This array is called SCALE because its elements should reflect the relative sizes of $(x_1, x_2, ..., x_n)$. Based on the required accuracy (ACC), the calculation finishes when, for $i = 1, 2, ..., n$, changes in xi of size ACC*SCALE(i) do not reduce the objective function.

### C Simulation methods

#### C.1 Coarse-grained MD

For all molecules, we ran a molecular dynamic in implicit solvent using Langevin thermostat with a time-step of 1 fs. Coarse-grained MD trajectories of 5 ns (during MD relaxation step in SAXS biased-method) or 150 ns (during MD SAXS unbiased-method) were performed at temperatures slightly below melting ($T \sim 100$ in our model's units), such that the molecule can rearrange easily but still aim at a folded structure.

Coarse-grained simulations for 1P5O, 1Y26 and 1A60 were performed at the experimentally reported ionic concentrations, corresponding to Debye-Huckel lengths of 1.8 Å, 3 Å and 4 Å respectively.

#### C.2 Atomistic MD

All-Atom molecular dynamic in explicit solvent were performed using Gromacs (version 5.0.7) [2]. Molecules were simulated in a cubic box of side of 1.6 nm filled with tip4p water. Monovalent and divalent ions ($K^{+}$, $CL^{-}$ and $Mg^{+2}$) were added to reproduce the physiological environment of biomolecules using molar concentrations of 0.15 M for KCl and 0.02 M for MgCl2. The system was then equilibrated in NVT ensemble using the velocity rescaling thermostat method, and then in NPT ensemble using Parrinello-Rahman barostat method [3, 4]. Temperature and pressure were fixed respectively at 300 K and 1 bar. Upon completion of the two equilibrations, an MD of 300 ns was launched. MD trajectory was then clustered using Gromacs cluster tool with 0.25 nm for RMSD cut-off parameter and gromos method [5].

### D Structure evolutions

## D.1 1P5O

| | Structure obtained after: | | | | |
|---|---|---|---|---|---|
| Starting conformation | 1st EM | 1st MD | 2nd EM | All-Atom MD | Native structure |
| 1-77 | 1-77 | 1-77 | 1-77 | 1-77 | 1-77 |
| | 2-76 | 2-76 | 2-76 | 2-76 | 2-76 |
| | 3-75 | 3-75 | 3-75 | 3-75 | 3-75 |
| | 4-74 | 4-74 | 4-74 | 4-74 | 4-74 |
| 5-73 | 5-73 | 5-73 | 5-73 | 5-73 | 5-73 |
| 7-72 | 7-72 | 7-72 | 7-72 | 7-72 | 7-72 |
| | 8-71 | 8-71 | 8-71 | 8-71 | 8-71 |
| 9-70 | 9-70 | 9-70 | 9-70 | 9-70 | 9-70 |
| 10-69 | 10-69 | 10-69 | 10-69 | 10-69 | 10-69 |
| | | 15-67 | 15-67 | | |
| 16-68 | 16-68 | 16-68 | 16-68 | 16-68 | 16-68 |
| 17-67 | 17-67 | 17-67 | 17-67 | 17-67 | 17-67 |
| | | | 18-64 | | |
| 18-66 | 18-66 | 18-66 | 18-66 | 18-66 | 18-66 |
| 19-65 | 19-65 | | 19-65 | 19-65 | 19-65 |
| 20-63 | 20-63 | 20-63 | 20-63 | 20-63 | 20-63 |
| | 21-61 | | | | |
| | | | | 21-62 | 21-62 |
| | | | 22-61 | 22-61 | 22-61 |
| | 22-59 | | | | |
| 22-60 | | | | | |
| | | 23-60 | 23-60 | 23-60 | 23-60 |
| 24-59 | | 24-69 | 24-59 | 24-59 | 24-59 |
| 25-58 | 25-58 | 25-58 | 25-58 | 25-58 | 25-58 |
| | 26-57 | 26-57 | 26-57 | 26-57 | 26-57 |
| 27-56 | 27-56 | 27-56 | 27-56 | 27-56 | 27-56 |
| | 28-55 | 28-55 | 28-55 | 28-55 | 28-55 |
| 29-54 | | 29-54 | 29-54 | 29-54 | 29-54 |
| | | | | 30-52 | |
| | | | | | 30-53 |
| | | | | 31-51 | 31-51 |
| | 31-50 | | | | |
| | | 32-50 | 32-50 | 32-50 | 32-50 |
| 33-49 | 33-49 | 33-49 | 33-49 | 33-49 | 33-49 |
| 34-48 | 34-48 | 34-48 | 34-48 | 34-48 | 34-48 |
| 35-47 | 35-47 | 35-47 | 35-47 | 35-47 | 35-47 |
| 36-46 | 36-46 | | 36-46 | 36-46 | 36-46 |
| 37-45 | 37-45 | 37-45 | 37-45 | 37-45 | 37-45 |
| | 38-44 | 38-44 | 38-44 | 38-44 | |
| | | | | | 39-43 |

Table 1: Evolution of base pairing (BP) for 1P5O. Green BP correspond to native, blue corresponds to BP off by one or two bases from native, red to BP off by more than two bases. Underlined are non-canonical BP. (EM: Energy minimization).

## D.2 1Y26

| | Structure obtained after: | | | | |
|---|---|---|---|---|---|
| Starting conformation | 1st EM | 1st MD | 2nd EM | All-Atom MD | Native structure |
| | | | 1-71 | 1-71 | 1-71 |
| | | | 2-70 | 2-70 | 2-70 |
| | | 3-69 | 3-69 | 3-69 | 3-69 |
| | | 4-68 | 4-68 | 4-68 | 4-68 |
| | 4-67 | | | | |
| | | 5-67 | 5-67 | 5-67 | 5-67 |
| | 6-66 | 6-66 | 6-66 | 6-66 | 6-66 |
| | 7-65 | 7-65 | 7-65 | 7-65 | 7-65 |
| | | | 7-64 | | |
| | 8-64 | 8-64 | | 8-64 | 8-64 |
| | | | | 9-63 | 9-63 |
| | 9-61 | | | | |
| | 9-62 | 9-62 | | | |
| | | | | | 10-40 |
| | | 10-61 | | | |
| | | | | 10-62 | |
| | | 11-60 | | | |
| | | | 12-59 | | |
| | | | | 12-42 | |
| | | 13-33 | 13-33 | | 13-33 |
| | | 14-32 | 14-32 | 14-32 | 14-32 |
| | 14-31 | | | | |
| | | 15-31 | 15-31 | 15-31 | 15-31 |
| 16-30 | 16-30 | 16-30 | 16-30 | 16-30 | 16-30 |
| | 17-29 | 17-29 | 17-29 | 17-29 | 17-29 |
| 18-28 | 18-28 | 18-28 | 18-28 | 18-28 | 18-28 |
| | | 19-27 | 19-27 | 19-27 | 19-27 |
| 20-27 | 20-27 | | | | |
| 21-26 | 21-26 | | | | |
| | 22-25 | | | | |
| | | | | 24-51 | |
| | 39-63 | 39-63 | 39-63 | | |
| 40-61 | | | | | |
| | 41-60 | 41-60 | 41-60 | 41-60 | |
| | | | | | 21-54 |
| | | | | | 22-53 |
| | | | | | 23-52 |
| | | | | | 25-49 |
| | | | | | 26-48 |
| | | | | | 34-41 |
| | | | | | 35-39 |
| | | | | | 42-60 |
| 43-59 | 43-59 | 43-59 | 43-59 | 43-59 | 43-59 |
| 44-58 | 44-58 | 44-58 | 44-58 | 44-58 | 44-58 |
| 45-57 | 45-57 | 45-57 | 45-57 | 45-57 | 45-57 |
| 46-56 | 46-56 | 46-56 | 46-56 | 46-56 | 46-56 |
| 47-55 | 47-55 | 47-55 | 47-55 | | 47-55 |
| | | | | 49-54 | |
| | | | | 50-53 | |

Table 2: Evolution of base pairing (BP) for 1Y26.

### D.3 1A60

In Figure 5 we report the three structures obtained by randomly modifying the orientations of the stems predicted by RNAfold, RNA Composer and Chimera for 1A60. For each initial structure we show the final structure obtained from the biased folding cycles. For S1 and S2 we show the structures chosen to be discarded, and for S3 the structure obtained at the end of the whole process (same as shown in the main text).

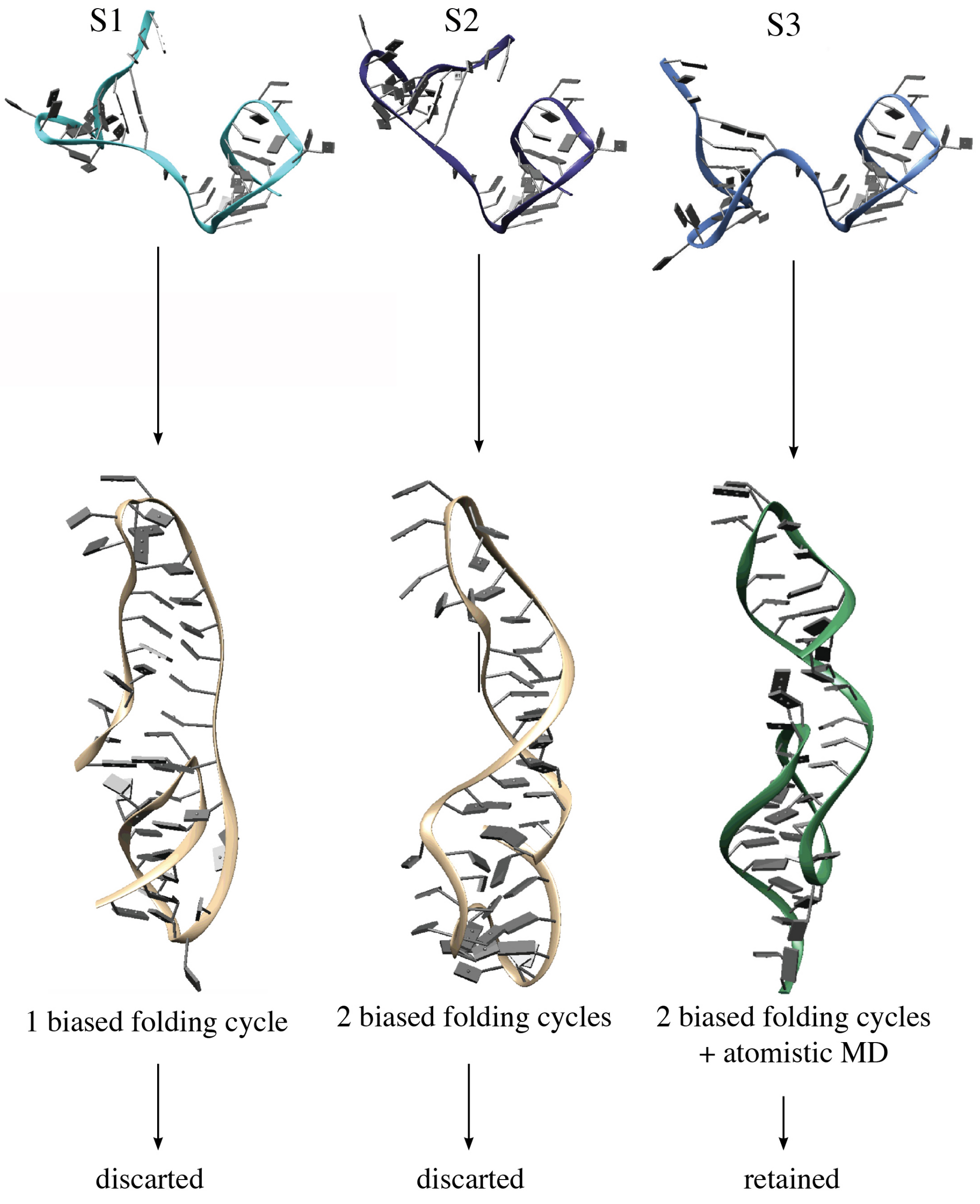

Figure 5: Initial structures used for fitting of 1A60 (top) and structures obtained from the biased folding procedure (bottom).

| | Structure obtained after: | | | | |
|---|---|---|---|---|---|
| Starting conformation | 1st EM | 1st MD | 2nd EM | All-Atom MD | Native structure |
| 1-17 | 1-17 | 1-17 | 1-17 | 1-17 | 1-17 |
| | | 1-18 | 1-18 | | |
| 2-16 | | 2-16 | 2-16 | 2-16 | 2-16 |
| | | 2-17 | 2-17 | | |
| 3-15 | 3-15 | 3-15 | 3-15 | 3-15 | 3-15 |
| 4-14 | 4-14 | 4-14 | 4-14 | 4-14 | 4-14 |
| 5-13 | 5-13 | | 5-13 | 5-13 | 5-13 |
| | | | | | <u>6-11</u> |
| | | | 6-12 | | |
| | | | 7-9 | | |
| | | | | | <u>7-10</u> |
| | | | | | 18-32 |
| | | | | | 19-31 |
| | | | | 20-30 | 20-30 |
| | | 21-30 | 21-30 | | |
| | 22-36 | 22-36 | 22-36 | | |
| | 24-40 | | | | |
| 24-41 | 24-41 | 24-41 | 24-41 | | 24-41 |
| | | | 24-43 | | |
| | | | | 24-44 | |
| 25-40 | 25-40 | 25-40 | 25-40 | 25-40 | 25-40 |
| 26-39 | 26-39 | 26-39 | 26-39 | 26-39 | 26-39 |
| | 27-38 | 27-38 | 27-38 | 27-38 | 27-38 |
| | 28-36 | 28-36 | 28-36 | | |
| | 28-37 | 28-37 | 28-37 | 28-37 | 28-37 |
| | | | | 29-36 | 29-36 |
| | | | | 30-34 | |
| | 33-35 | | | | |

Table 3: Evolution of base pairing (BP) for 1A60.

### E Comparison with conventional fitting method

We present here the comparison of our results with the widespread approach of fitting a structure into an envelope computed from an intensity curve $I(q)$. For structures which must undergo a significant deformation from the initial configuration to achieve a fit (as it is the case for partially unfolded molecules), this can be done using the tools provided in the ATSAS software [6] combined with the program iMODFIT, available as a plugin for Chimera visualization software [7]. iMODFIT performs normal modes in internal coordinates, and it is therefore capable of substantially modifying a structure while fitting. Following what can be done with these tools, we first compute an envelope and perform a fit of the partially unfolded structures using iMODFIT. This brings the overall shape of the molecule to approach SAXS data but it does not rearrange local structures. To achieve local structure formation, we perform a short atomistic MD simulation of 10 ns and we extract the final structure to perform a second fit, using Chimera volume fit function, which performs only translations and rotations, therefore not altering the internal structure obtained in the MD.

#### E.1 SAXS "envelopes" and fitting method

DAMMIF program from ATSAS software [8] was used to compute envelopes from scattering curves, as strongly recommended for nucleic acids [9]. The scattering curves used to generate the envelope for the native (target) structures were computed from Debye theory at atomistic resolution in vacuo. The program GNOM was used to generate the input files required by DAMMIF from the scattering curve. It requires as inputs, the intensity curves and a maximum particle diameter $R_{max}$. As indicated by default, $R_{max}$ is taken as 3 times the radius of gyration of the molecule. In our case we set $R_{max}$ 90 Å for 1P5O, and 65 Å for 1Y26 and for 1A60.

The program iMODFIT, was used in Chimera [7], to fit structures into the envelopes using normal modes analysis. This allows to perform more significant transformation in the structure compared to the programs provided in ATSAS that perform only rotations and translations of the whole molecule. iMODFIT has been used for flexible fitting of structures into EM maps based on Normal Mode Analysis in Internal Coordinates [10], and it has been used for proteins. iMODFIT default parameters were used.

A second fit to the envelope was performed using Chimera volume fitting tool (translation and rotations only) after a 10 ns MD all atom simulation to relax the structure obtained by iMODFIT in order to stabilize local structures and form base pairs.

#### E.2 Fitting results

A summary of fitting results is presented in Figure 6.
For 1P5O, at the end of this fitting procedure we obtain a structure with RMSD of 8.3 Å from the native, with a fitting score $\chi = 2.7 \times 10^{-6}$, 26 base pairs, 23 of which corresponding to native (out of 32) 3 and off by one or two bases. For 1Y26, we obtain a structure with RMSD of 15.2 Å from native, $\chi = 8.9 \times 10^{-6}$, and 18 base pairs, of which 5 native, 10 off by one or two bases, 3 off by more than two bases. The kissing loop is not formed and if one performs a simulation starting from this structure without any constraint the structure opens and the two inner stems drift apart. For 1A60, the final structure has an RMDS of 14.9 Å from native and $\chi = 9.0 \times 10^{-6}$. It forms 11 base pairs, of which 8 native, 1 off by one base and 2 off by more than two bases. The pseudoknot is not recovered, and the structure does not appear to be stable (that is a longer dynamics results in a significant drift away from the fitted structure).

The time required to complete this fit for each molecule was of about dozen hours, using 10 hrs of high performance parallel computing with around 252 cores.

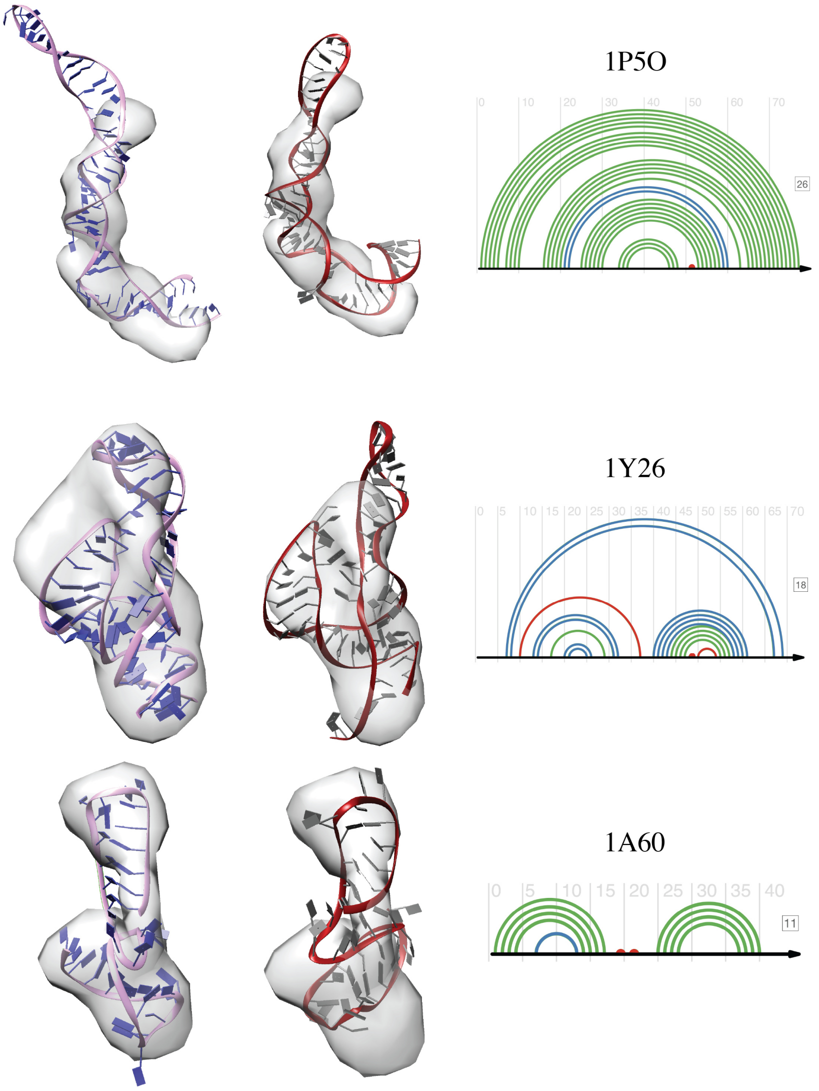

Figure 6: Results of fitting procedure using DAMMIF and iMODFIT. For all systems, left (pink): structure resulting from fitting the partially unfolded structures into the envelope calculated by DAMMIF; center (red): structures extracted via a short atomistic MD are fitted into the envelope; right: base pairings of the structure obtained from the atomistic MD.
